# *Virotest*: a bioinformatics pipeline for virus identification in plants

**DOI:** 10.1101/2023.02.15.528703

**Authors:** Daniel Gonzalez-Ibeas

**Affiliations:** personal

**Keywords:** plant, crops, virus, bioinformatics, deep sequencing, small RNAs

## Abstract

High-throughput DNA sequencing is being introduced as an emerging technology for viral diagnosis. Despite several tools being available, lack of standardized bioinformatic pipelines and price are major limitations for its implementation. Many extant tools focus on the assembly of virus-derived small RNAs into contigs and their subsequent annotation. Better sensitivity has been reported by several studies when this approach is complemented with read mapping on known viral genome sequences as an ancillary step, requiring less number of reads and allowing price optimization during DNA sequencing. This publication introduces *virotest*, a bioinformatics pipeline for the identification of viruses in plants from small RNA data. The tool prioritizes read mapping over sequence assembly for the purpose of obtaining better sensitivities and requiring lower sequencing depths.

## Introduction

Plant viruses are important pathogens that cause severe losses in agriculture [1], [2], and almost 50% of the microorganisms responsible for emerging plant diseases are viruses [3], [4]. Along with cultural practices (such as vector management or weed control), and immunization (mostly driven by genetic resistance), early detection of viruses is part of the main strategies for disease control, and it is frequently employed to propagate virus-free plant material or to monitor virus prevalence [5]. Nowadays, globalization favors viral dispersion across countries due to migratory movements and plant material exchange, in such a way that introduction of invasive species is increasing and expected to grow [6]. Climate change is compounding the problem of viral epidemics, disease outbreaks and global food security [7]–[9]. Efforts have been started more than 10 years ago to establish general guidelines for plant material quarantine, virus identification and naming rules of new species or strains in the current context [10], [11].

Enzyme-linked immunoassay (ELISA) has been over decades one of the most popular conventional methods for virus detection [5], [12], [13]. However, it lacks the resolution to properly identify some viral strains. On the contrary, molecular probes based on nucleic acids, such as dot-blot, tissue print, polymerase chain reaction (PCR), real-time quantitative PCR (qPCR) or isothermal PCR, can be generated at low cost and have the required flexibility to deal with the distinction of viral sequence variants that do not rely on amino-acid changes [14]–[17]. Multiplexing and high-throughput techniques have been also applied to virus detection, such as microarrays, but some limitations persist related to cross-hybridization and the lack of capabilities to discover uncharacterized viruses.

As an alternative to conventional molecular methods, next-generation DNA sequencing (NGS) is being introduced as an emerging technology for diagnosis [5], [18]. A larger investment in molecular biology and bioinformatics expertise makes this technology only suitable for well-funded and centralized laboratories, according to some authors [13]. However, this situation is not new and the same criteria once applied to techniques such as qPCR, and over time, the cost of sequencing has dropped at an incredible rate and several analysts envision a price per sample similar to ELISA or PCR tests [12], [13]. NGS technology offers the opportunity to detect new viruses not previously reported on databases [13], since no probe design is required. Additionally, the collection of nucleic acid samples from the environment allows circumventing culture steps to capture the whole diversity in the sample. This is of particular interest with regard to viruses, because 1) of their abundance as biological agents on Earth [19], [20], 2) they are obligate parasites that need a host to replicate and cannot be grown on artificial culture media, 3) co-infections are frequent in plants leading to several types of effects, such as amplification/attenuation of symptoms, or promoting recombination events which may be behind the emergence of new strains. Moreover, the beneficial role of bacteria or fungi in ecosystems as symbionts is well-known long ago, but this is not the case for viruses, and investigations have focused almost exclusively on their pathogenic role. However, current studies are suggesting that viruses could also live in symbiosis with their hosts, being commensals and providing a neutral balance to the relationship, or even providing a benefit in the interaction with bacteria, insects, plants, fungi and animals [21]–[24]. As a consequence, non-pathogenic viruses may co-exist with crop-infecting species and might contribute to all processes that viruses have been associated with. In the results section, a potential example in rice of such co-existence is shown. Sometimes the identification of commensal viruses will interfere and be background noise for the method, and other times will be informative for species novelty and pathogenesis, but these considerations are relevant for virus diagnosis and NGS proves an opportunity to address them.

For the development of a such diagnostic tool, major challenges come from the lack of gold standards in the methodology, especially about bioinformatic tools [12], the type of viral molecule to be screened, and the high cost of the analysis to be applied routinely. Virus-derived small RNA (viRNA) identification has become a popular choice for diagnosis. For novel viruses characterization, *de novo* assembly of the viral genome is often the preferred approach, and high sequencing depth is performed to catch the whole viRNA population, which makes price optimization difficult. In comparative studies of methodologies, mapping read data in combination with assembly notoriously improves sensitivity [25], and other studies have reported that when the viral genome sequence is known, several orders of magnitude of fewer reads are required for proper detection by mapping-based analyses, reaching a sensitivity similar to qPCR [26]. Thus, prioritizing read mapping over sequence assembly could offer an opportunity to reduce the required sequencing depth, and also to avoid the issues associated with viral contig assembly from sRNA data of low-titer viruses [27]. Several computational tools are currently developed that combine and/or prioritize, to varying degrees, both read mapping and assembly of contigs (see [28] for a review). This publication introduces *virotest*, a bioinformatic pipeline to detect viruses in crops from small RNA data based on read mapping significativity with a main emphasis on economical cost and sensitivity. Simulated and real data downloaded from repositories have been used to assess the performance of the pipeline.

### Molecular approach

The analysis of double-stranded (ds)-RNA, viral particles (virions) or viRNAs is usually employed in viral metagenomics. All of them have advantages and disadvantages to be used for diagnosis, and none of them have been set as “universal” to capture the real diversity of viruses. Thus, 1) (ds)-RNA isolation is based on the fact that endogenous plant RNAs do not form extensive double-stranded structures, contrary to what replicative intermediates of RNA viruses do. However, among RNA viruses, this method is biased towards ds-RNA viruses [29], it is not suitable for the identification of some DNA viruses (because little or no dsRNA is generated), and it also fails detection of persistent (cryptic) viruses, which can stay as “silent” without undergo multiplication [29]. 2) Virion purification has been used for DNA virus screenings [30], [31] and cryptic viruses, but fails the identification of viroids, important causal agents of disease in some crops [32]. 3) Small RNA sequencing is used to detect viRNAs generated as part of the immune response of the plant based on gene silencing. The method is effective for the detection of viroids, RNA- and DNA-viruses, but it fails identification of cryptic viruses. Although the best approach to capture the whole viral diversity is perhaps a combination of all techniques, sRNA sequencing might be of particular interest for diagnosis because of two reasons: first, while ds-RNA or virion purification methods are time-consuming, experimentally challenging and rely on sophisticated and expensive laboratory equipment (e.g. ultracentrifugation), small RNA purification is becoming a more straightforward task; second, known plant pathogenic viruses undergo cellular multiplication and subsequent viRNAs biogenesis, whereas cryptic viruses tend to be non-pathogenic, thereby failure in their detection from sRNA data could not be a major problem. As a consequence, sRNA purification, sequencing and viRNA identification is gaining in popularity and has been successfully used for *de novo* assembly of viral genomes [33], [34]. Two sources of contamination are worth considering for the proper use of viRNAs as a diagnostic tool:

1. Fungi can produce sRNAs as part of their metabolism that can be co-purified with plant-derived molecules if the sample is contaminated with either symbionts or pathogenic fungi, not just because of the presence of hyphae in the plant material, but also because the movement of sRNAs between plants and fungi has been documented [35]. This may be of special relevance if the fungus is a biological vector of plant viruses [36], or if mycoviruses (and their associated sRNAs) are present in the fungus [37].
2. Plant genomes are populated with regions of presumed viral origin, most of them with similarity to DNA viruses, but in rare cases, segments from RNA viruses have been also reported [38]–[41]. There exists the possibility that these virus-like areas become a source of sRNA with similarity to viral genomes [42]–[45], representing an important source of confusion for viRNAs categorization.

### Bioinformatic pipeline implementation

The pipeline starts with a first step of sanity check and statistics of the input file in FASTA format. Next, the pipeline subtracts sRNAs mapped on the host plant genome to reduce noise and increase the efficiency of pathogen detection [46], [47]. Next, the pipe has two parts (Figure 1). Step 1: Read mapping on a custom-made database using a non-redundant combination of viral genome sequences from the DPV (http://www.dpvweb.net/) and GenBank Viral RefSeq (http://www.q-bank.eu/Virus/) databases (totaling 1,131 viral entities). Genome sequences with more than 95% of identity are clustered as one single representative sequence. Only plant viruses are used to improve runtime and mapping efficiency, as suggested [46]. Read mapping is performed with micro-razers from the SeqAn library [48]. Known viruses potentially present in the sample are identified using interquartile range (IQR) to detect outliers in mapping rates with the *Scipy* [49] and *Numpy* [50] Python libraries. Five parameters are implemented and combined for developing purposes, including the percentage of sRNAs with a hit on the viral genome sequence (T1), percentage of the viral genome covered by uniquely mapped sRNAs (i.e., non-mapped on other viral genomes of the database) (T2) or by sRNAs mapped on several entries of the database (T3), uniqueness (percentage of T3 due to T2) (T4), and percentage of sRNAs (from the total sRNA read set) uniquely mapped on the genome (T5). IQR performance requires additional filtering to avoid false positives, but it allows complexity and runtime reduction in the calculation steps. Identified viral genomes by the pipeline are classified as “high” or “low” quality, the former as highly probable viruses in the sample, the latter as inconclusive output [11]. Both results are browsable to check up on ambiguous classifications. An output example of the pipeline is depicted in Supplementary Figure S1. Step 2: sRNA reads are assembled into contigs with Velvet [51]. Assembled contigs are queried against the custom database to re-evaluate results from the mapping analysis. Inconsistencies in the comparison are used to detect a potential “novelty” in the sample. Additionally, remote BLAST analysis of unmapped contigs against the non-redundant (nr) GenBank database is performed with the NCBIWWW module of *Biopython* [52]. The pipeline is deployed as a website with Apache (https://httpd.apache.org/) and PHP (https://www.php.net/) on an Amazon Web Services (AWS) cloud, which can be set up upon request. It runs under a Linux environment (kernel 5.18.10-76051810-generic) and all necessary software in integrated using Python as a wrapper with the *subprocess* module.

**Figure 1.**
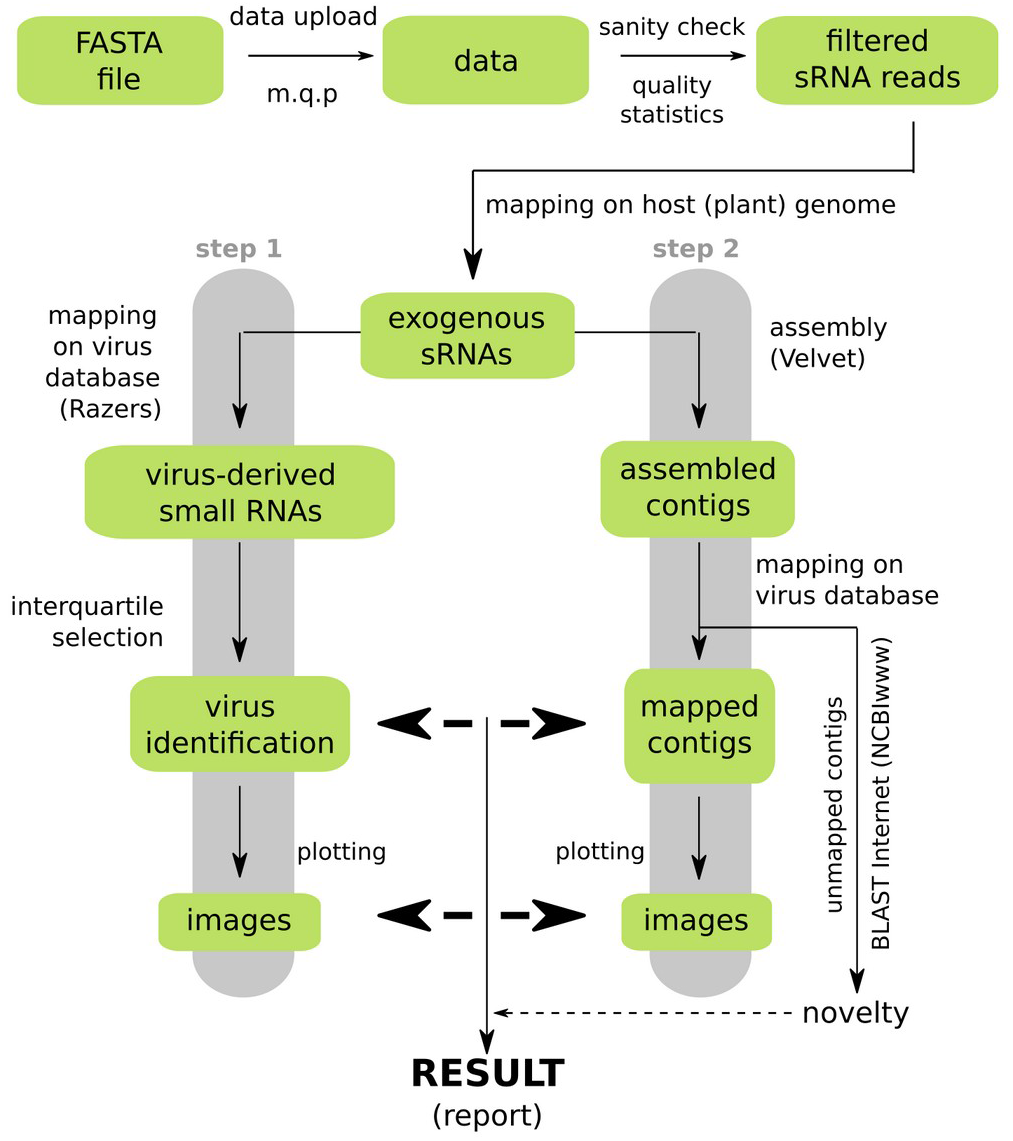
Flowchart of the *virotest* pipeline.

## Results and discussion

Calculation of the theoretical sensitivity (i.e, minimum number of reads for a virus to be significantly identified) could help to optimize the pipeline (based on the virus-crop target) or the experimental procedure (for example the number of Illumina libraries per lane, or to escalate a pooling strategy when several plants are pooled from the same cropland). Artificial small RNA reads have been generated with custom Python scripts from the viral genome sequence of every entry in the custom database, as a proxy of real viRNAS to estimate sensitivity and specificity [11]. The data have been grouped into 10 read sets ranging from 11 to 6,000 reads. Each read-set for each virus has been used as input for the *virotest* pipeline, and the number of times the virus was identified as “correct” and “unique” was counted. Even with only 93 reads, 98% of the viruses could be properly identified using the mapping approach alone (with no assembly) (Figure 2). A positive correlation between the length of the viral genome and the minimum number of reads (based on parameter T3) was visible (Figure 2), as expected. Even just 6,000 reads seem enough for proper genome coverage and sensitivity for large viruses.

**Figure 2.**
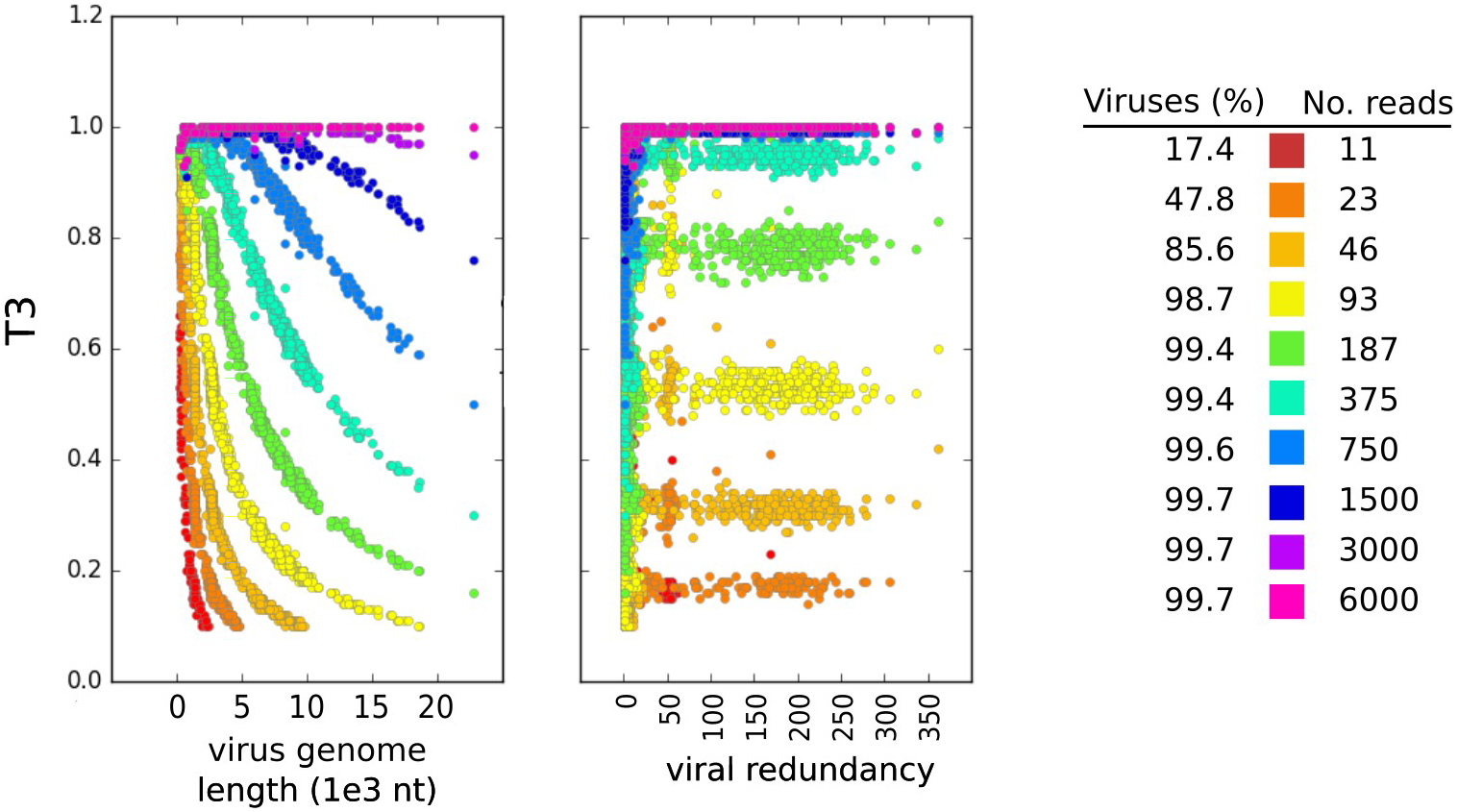
Analysis of T3 parameter results. Each dot represents the resulting T3 value from virotest for each viral accession of the custom database during the analysis of simulated data. T3 is defined in the pipeline as the percentage of the viral genome covered by sRNAs. T3 values were plotted against the virus genome length (A) and the redundancy level for each virus (B), defined as the number of viruses that shared sequence similarity at 70% genome coverage and 70% identity. The number of simulated reads from the genome sequence of each virus used in each iteration is shown in the legend, along with the percentage of viruses (of the custom database) identified as “unique” and “high-quality” by the pipeline.

The representation of viral families in the custom database was highly skewed (Figure 3). The most abundant were *Geminiviridae, Potyviridae* and *Betaflexiviridae*. Although sequence redundancy has been decreased at the level of 95% of identity during the custom database creation, the influence of the sequence similarity among viral entries might have an impact on the performance of the pipeline, since the proportion of uniquely mapped or shared viRNAs is used to detect positives in the samples, similar to other approaches [53]. Thus, the addition of several closely related viral isolates to the database (instead of using representative viral species) is worth optimization for this pipeline. The length of genome areas of each virus with sequence similarity to other members of the database was used as an estimation of the virus redundancy rate. When the read-sets were analyzed with respect to this parameter, as opposed to the viral genome length, no glaring correlation was seen (Figure 2). The average genome coverage was affected by the depth of the sequencing (as expected) but it was evenly distributed for every redundancy rate. The departure of some values from this trend (Figure 2) was manually reviewed and the large genome size seemed the main cause.

**Figure 3.**
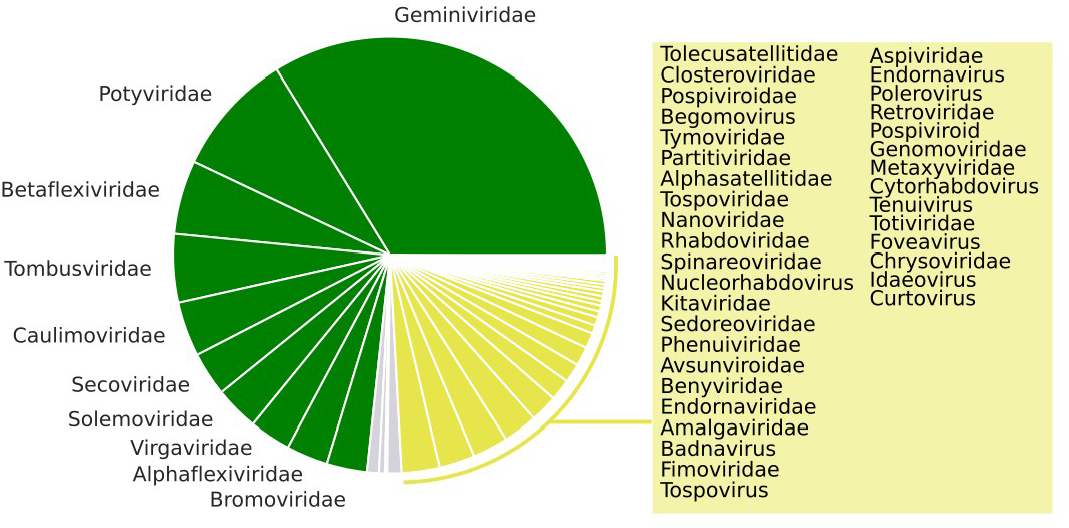
Taxonomic composition of the custom database by family. The figure depicts high (green) and low (less than 3%, light green) abundance taxa. The latter is collapsed to improve readability. Unassigned entries are shown in grey.

In real data, the proportion of viRNAs from the total plant sRNAs may vary depending on several factors, including the virus accumulation rate, the stage of the infection, the plant tissue, or the host-virus combination at the species level, among others. Thus, sRNA sequencing data downloaded from public repositories were used to evaluate the performance of the pipeline (Table 1). Included are 10 samples from 3 crop species that have been used for a broad comparison of different workflows for virus diagnosis in plants [25]. This study has been self-proposed as a benchmark for pipeline development. Each sample represents a subsample of reads simulating 3 sequencing depths (i.e., 50K, 250K and 2,500K reads). During the *virotest* assessment, the most relevant result was the identification of *Apple stem grooving virus* (ASGV) in the apple samples at the 250K and 2,500K depths. In the reference study, those samples were excluded because too few participants successfully reported the identification of viral contigs [25]. This result suggests that a mapping strategy might overcome some of the problems of sensitivity reported by other approaches. *Vitis* samples were reported of particular difficulty since the plants were infected with at least 9 viral species, confirmed by molecular methods [25]. All of them were identified by *virotest* at 2,500K depth (Table 1), but some failed when fewer input reads were used. For *Grapevine rupestris stem pitting-associated virus* (GRSPaV) and *Grapevine leafroll-associated virus* (GLRaV), two additional strains were identified as “low-quality” and not considered for the main results, since both showed non-significant mapping rates of uniquely mapped viRNAs (Figure 4). In the case of GLRaV, both strains show 90% of identity over more than 90% of the genome, revealing that this identity level allows distinction between viral strains by mapping rates given the current threshold (95%) used to reduce redundancy in the custom database. This parameter could be tuned up, if needed. The pipeline automatically prioritizes uniquely mapped viRNAs to select the most probable accession in the sample. In the case of GRSPaV, an additional strain was also identified as “high-quality”. The abundance of unique viRNAs suggests that both strains of this virus might be present in the sample. For every result, all high- and low-quality accessions are browsable (see Supplementary Figure S1) to allow open interpretation by the pipeline’s end user.

**Figure 4.**
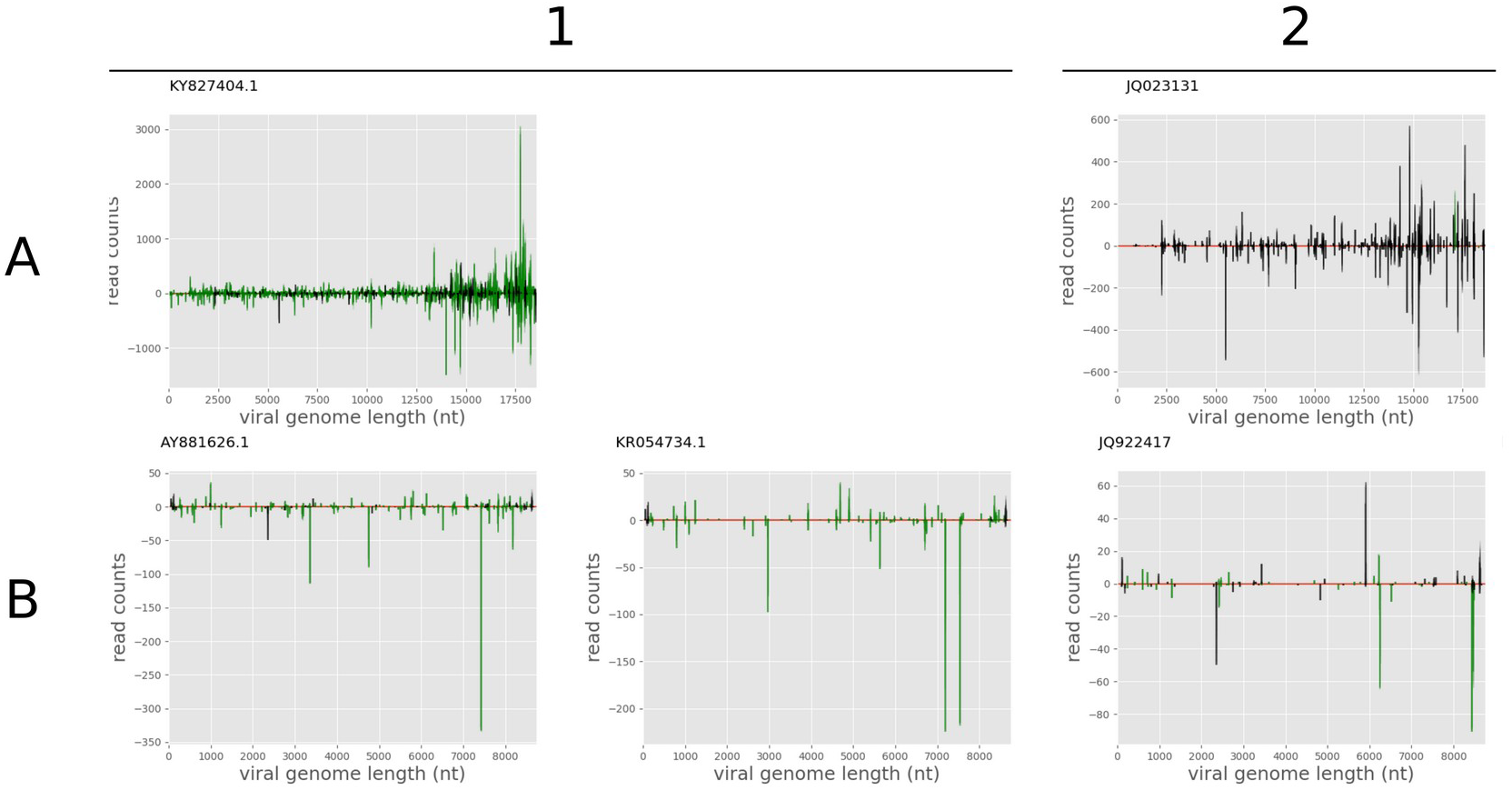
Number of sRNAs mapped on each nucleitide position of the viral genome of *Grapevine leafroll-associated virus* (A) and *Rupestris stem pitting-associated virus* (B), from Sample_6 of Table 1. Results are shown for viral strains identified as “high-quality” (1) or “low-quality” (2) by *virotest*. For each plot, counts of sRNAs uniquely mapped (green) or shared with other viral entries of the database (black) are shown. The viral genome is represented by a red line. Read counts correspond to forward (positive values) and reverse (negative values) sRNAs.

**Table 1.**
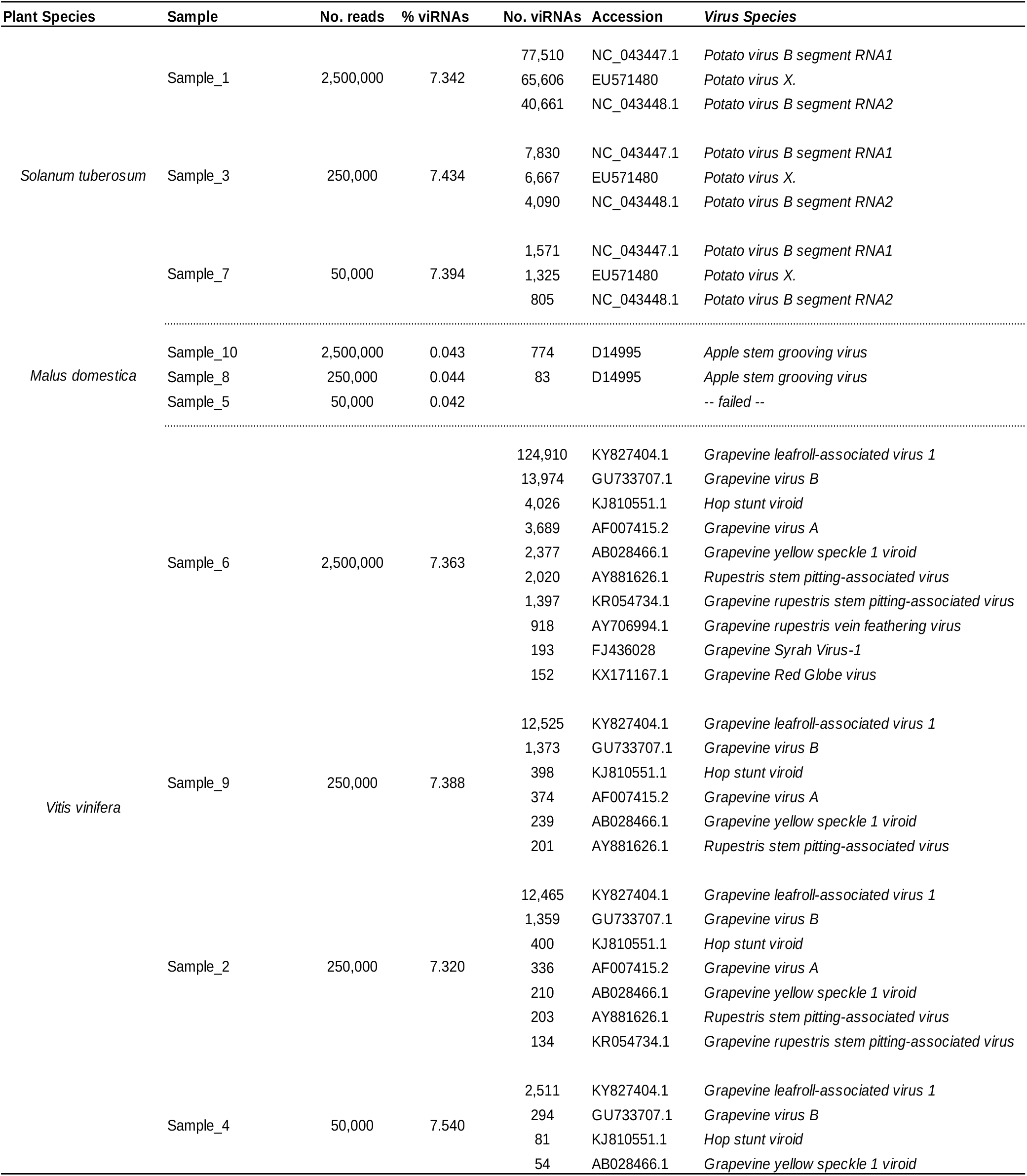
Viruses identified bv virotest in samples from a reference study for pipeline development.

In addition to the reference study, other sequencing data were included in the evaluation (Table 2). Raw files were subsampled to simulate the same depths as for Massart and others [25], for consistency. In the rice samples, *Rice stripe virus* (RSV) is properly identified at 250K depth in infected samples in agreement with the published work. Additionally, *Oryza sativa endornavirus*, seemingly not reported in the publication, was also found. Plant endornaviruses have several persistent (symbiotic) properties compared to other conventional viruses, including a stable low copy number, no obvious effect on the host plant, and efficient vertical transmission via gametes (they are likely maintained within host plants for hundreds of generations) [54], [55]. Indeed, virus-carrier and virus-free plants are reportedly indistinguishable in some rice cultivars [56]. RSV was identified in the source publication by ELISA and viRNA mapping only on the RSV genome [57]. The reported findings by that work are relevant, of high quality and valid, but the *virotest* result presented here exemplifies the benefits of unsupervised methods in the detection of viruses. The presence of this additional endornavirus was confirmed by VirusDetect [46]. Surprisingly, this software failed the identification of RSV at 250K depth, and 2,500K reads were required, suggesting again that prioritizing a mapping approach above the assembly might provide higher sensitivity. Other cases where *virotest* findings agreed with those reported in the source publications are depicted for peach, maize, cotton, papaya and tomato (Table 2). The peach samples revealed infection by several viruses and proper identification required the highest depth. When 250K was used, *Plum pox virus* failed identification in sample PEACH.GRR.PPV, and at 50K, detection of *Prune dwarf virus* and *Peach associated luteovirus* also failed in several samples (a full breakdown is shown in Supplementary Table S1).

**Table 2.**
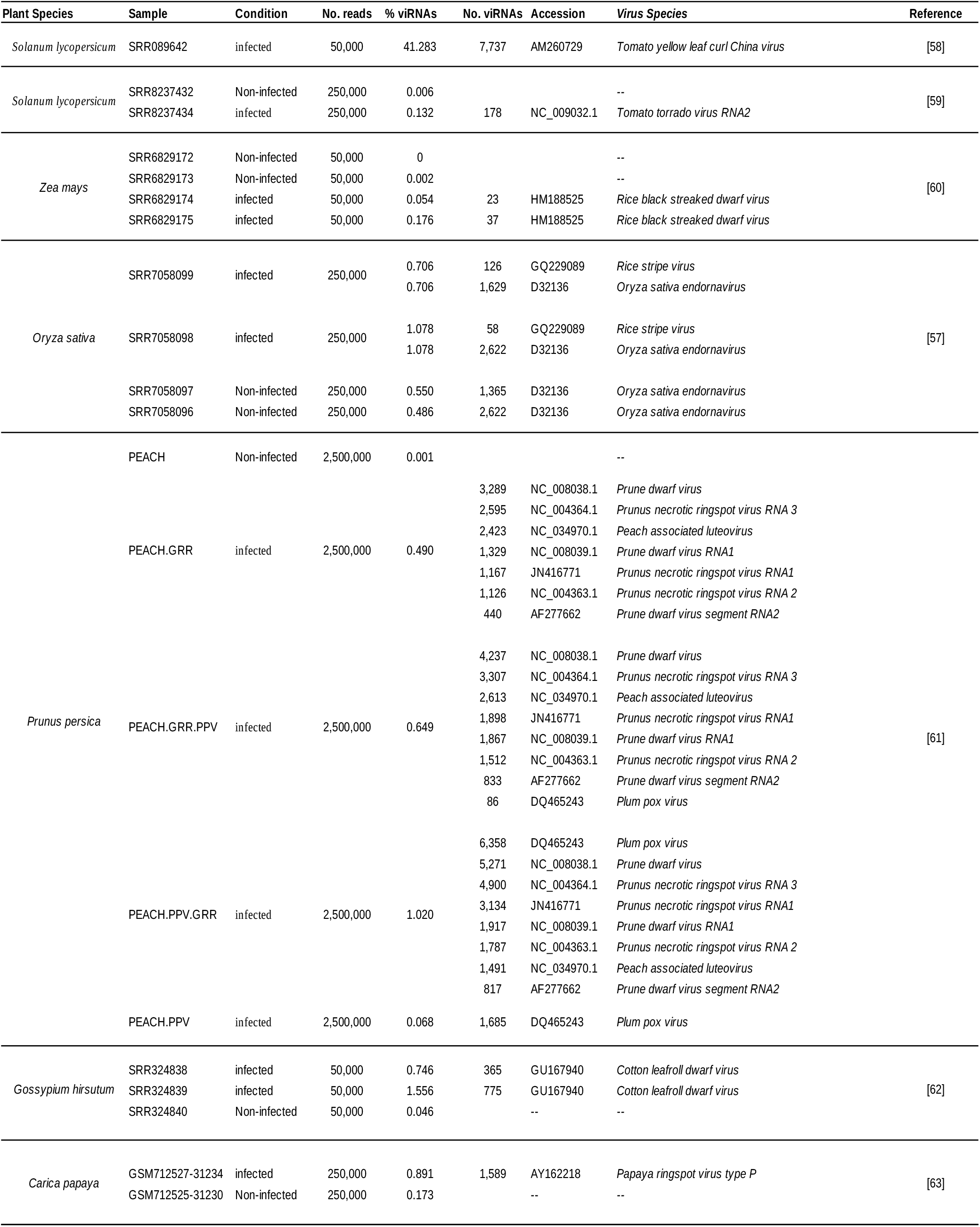
Viruses identified in several crop species by the virotest pipeline in samples from published studies.

The use of short sequence data comes with a risk of including reads that overlap with viral genome areas conserved among viral species, leading to false positives. Thus, several cases were reported by different pipelines in the reference study, including the identification of species of the *Caulimoviridae* family in potato and grape samples [25]. Seemingly, this was not a problem in the *virotest* pipeline since no additional viruses were detected when compared to those reported in the source publications. Lastly, the pipe implements a novelty factor to detect cases of viruses absent in the custom database. *Vitis* samples (Table 1) are an example, which showed the highest levels of discrepancy in the comparison between mapping and assembly rates of the pipeline due to the absence of specific strains (data not shown). After the manual addition of these virus accessions to the custom database, the novelty factor dropped. These cases can be quickly identified and corrected by browsing the BLAST results of unmapped contigs in the *virotest* output, as well as cases where potential new viruses (or isolates) have similarities to other known viruses. However, if the novel viruses were phylogenetically distant, the same limitations of other homology-based approaches would apply here [25], [46].

## Conclusions

In summary, although a generalized depth of 2,500K has been proposed [25], viruses from many samples presented in Tables 1 and 2 are identifiable at lower depths, representing more than one order of magnitude for optimization. The results of AGSV in apple and RSV in rice suggest that prioritizing the mapping approach could be an improvement over assembly-based methods. However, the results of *virotest* in grape and peach reveal that higher depths might be required, at least when low viral titers or co-infections are expected. Further development could include customization of the composition of the database or the selection parameters used here, in order to tune up the pipeline to tailor specific viruses or crops (e.g. woody vs non-woody plants would be an option). In any case, the minimum number of reads required for proper diagnosis in real samples is far above the theoretical values for computational significance analyzed here, due to the low percentage of viRNAs in plant sRNAs. In this regard, sequence capture technologies could provide a solution for viRNA enrichment, but in price-centered diagnosis, this probably would make the method economically unaffordable. The main drawback of the approach presented here is that extremely low sequencing depths (to optimize cost) play against the proper assembly of viral sequences. Generally speaking, the assembly of contigs is probably an essential step when novel viruses are expected (or suspected) but, yet virus emergence is an actual threat in agriculture, there is a sweet spot between the incidence of emergence events and the number of samples to be analyzed in day to day large scale screenings, especially when detection of known viruses requires economical affordability for routine analyses.

