## Supplemental Figure 1 for "*Virotest*: a bioinformatics pipeline for virus identification in plants"

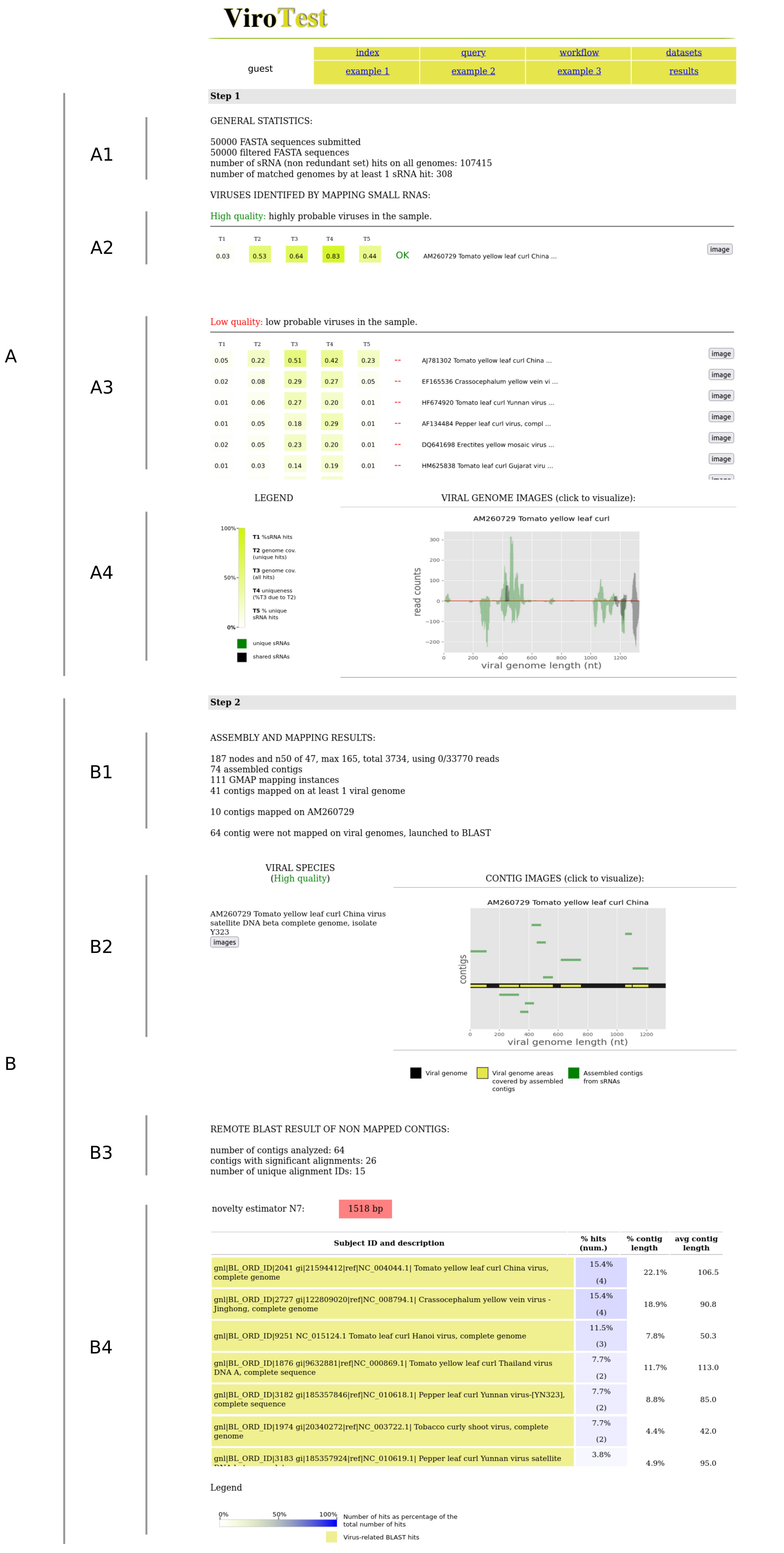


Supplementary Figure S1. Output example of the virotest pipeline corresponding to accession SRR089642 from Table 2 of the main text. Step 1 of the pipeline: sRNA mapping on the custom viral genome database (A). Input read data and mapping statistics (A1). Virus accessions identified as “high-quality” (A2) or “low-quality” (A3) and the five ratios (T1-T5) used for filtering. Plot of number of sRNAs uniquely mapped (green) or shared with other viral entries of the database (black) on each nucleotide position of the viral genome (A4). Legend of the five rations (T1-T5) is shown here in green scale color (from 0 to 100%) to make interpretation easier. Step 2 of the pipeline: sRNA assembly and BLAST results (B). Assembly statistics (B1). Plot of contigs mapped back on the genome sequence of viruses identified as “high-quality” in the sample by step 1 (B2). BLAST result statistics of unmapped contigs against the non-redundant nucleotide database of GenBank (B3). Non-redundant accession IDs from BLAST results in decreasing order sorted by the number of hits and colored in a blue scale (for readability) (B4). The number of assembled nucleotides (from the total number of contigs) with no BLAST hits is shown in red. Virus or viroid-like BLAST descriptions are colored in green background.
